# LuTuFlow: a deep learning-based pipeline for quantitative characterization of mouse lung tumors in CT imaging

**DOI:** 10.64898/2026.10.12.758509

**Authors:** Yuchen Liu, Luqing Li, Mallory Wittwer, Amaia Martinez-Usatorre, Nahal Mansouri, Bastien Dapia, Edward Andò, Michele De Palma

**Affiliations:** Swiss Institute for Experimental Cancer Research (ISREC), School of Life Sciences, Swiss Federal Institute of Technology in Lausanne (EPFL), Lausanne, Switzerland; Agora Cancer Research Center, Lausanne, Switzerland; Swiss Cancer Center Léman (SCCL), Lausanne, Switzerland; Center for Imaging, EPFL, Lausanne, Switzerland; Division of Pulmonary Medicine, Department of Medicine, Lausanne University Hospital (CHUV), University of Lausanne (UNIL), Lausanne, Switzerland

## Abstract

Computed tomography (CT) is a widely used, non-invasive imaging tool to monitor tumor growth. Tumor identification and volumetric quantification are crucial for assessing tumor burden, whereas tracking individual tumors by aligning scans from multiple time points reveals disease progression. However, these processes remain largely dependent on manual analysis, limiting the accuracy, reproducibility, objectivity, and scalability of tumor characterization. Several deep learning models have enabled automated segmentation and object identification. However, open-source, integrated software platforms that cover the entire workflow – including processing raw CT images, annotating and tracking tumors through time, and generating data reports and visualizations – are still needed. Here, we present LuTuFlow, a computer vision pipeline incorporating a nnU-Net-based deep learning model, configured for murine lung segmentation and tumor analysis. We integrated these modules into a Napari plugin, allowing researchers to apply the entire automated workflow to transform raw CT scan datasets into quantitative readouts and interactive visualizations in a simple graphical user interface. The resulting model, iteratively refined using expert-corrected annotations, generated reliable quantitative estimates of tumor burden.

## Introduction

Lung cancer is among the most prevalent cancers and is the leading cause of cancer-related mortality worldwide^1^. Mouse models of lung cancer have been developed that recapitulate key features of human disease. Among these, a genetically engineered mouse model (GEMM) with lung-specific, conditional mutations in *Kras* (leading to KRAS hyperactivation) and *Trp53* (leading to p53 loss of function), commonly termed the “KP” model, recapitulates the histological and molecular features of a subset of human non-small-cell lung cancers (NSCLCs)^2,3^. These mice develop lung tumors over several weeks, with significant variation in the kinetics of disease onset and progression^2,3^. KP cancer cell lines isolated from this GEMM can also be used to establish synchronous, orthotopic lung tumors in wild-type recipients^3^. In this setting, the resulting tumors may lack the genetic and histological heterogeneity of tumors genetically induced in the GEMM^3^.

Spatial localization and volumetric size are two crucial readouts for evaluating tumor growth in preclinical oncology studies. Computed tomography (CT) provides a non-invasive, high-resolution tool for *in vivo* imaging of disease sites^4–7^. Individual tumors can be identified and measured from CT scan images, and scans from multiple timepoints enable tumor tracking for quantification of tumor progression. These tasks have traditionally relied on manual tumor identification and tracking, with software applications such as RadiAnt DICOM Viewer^8^, ITK-SNAP^9^, OsiriX^10^, and AnalyzeDirect^11^ providing robust functions. However, manual assessment is susceptible to human error and inter-observer variability, which may limit the accuracy, reproducibility, and objectivity of tumor analyses^6,7^. Moreover, such an approach has limited scale-up potential and can be time-consuming when analyzing datasets with large numbers of mice or tumor nodules.

To address these challenges, several automated image analysis tools have been developed to streamline CT scan interpretation and tumor identification. These include deep learning (DL) models designed for segmenting the lung parenchyma in biomedical images^12^, as well as for volumetric segmentation of lung cancer in both human^13,14^ and mouse^15,16^ CT images. Despite these advances, open-source, integrated software platforms that cover the entire analysis workflow (including processing raw CT images, annotating tumors and tracking them through time, and reporting on indicators that reflect disease progression) are still needed.

Modern platforms like Ultralytics YOLO^17,18^ and nnU-Net^19^ enable implementing computer vision (CV) pipelines that offer high accuracy and robustness, without requiring the design of custom model architectures or training recipes. In particular, nnU-Net^19^ provides self-configuring 3D segmentation models as well as a standardized machine learning workflow that can be used to efficiently train specialized models capable of addressing the segmentation tasks encountered by individual research groups. Tracking individual tumor nodules can be achieved by performing an initial spatial alignment of the scans and then matching individual tumors across scans based on their position and appearance, using tracking algorithms such as those implemented in the LapTrack^20^ library. Additionally, CT images and outputs from the CV pipeline, including segmentation masks and individual tumor tracks, can be visualized, inspected, and edited interactively in Napari^21^, an open-source n-dimensional image viewer based on Python. Napari also offers a graphical user interface that can be extended with plugins, allowing the development of customized platforms for tumor characterization from biomedical images.

Here, we present LuTuFlow, a CV pipeline incorporating a nnU-Net–based DL model, tailored to murine lung segmentation, tumor identification, and tumor tracking in sequential scans. We integrated these modules into a Napari plugin, allowing researchers to access the workflow from a simple user interface (UI). Tumor identification performance was iteratively improved through expert correction of model predictions, and the resulting model demonstrated robust detection of lung tumor nodules with reliable quantitative accuracy.

## Methods

### Mouse models of NSCLC

The *Kras*^LSL-G12D/+^;*Trp53*^fl/fl^ (KP) transgenic mouse model (C57BL/6 genetic background) of NSCLC was maintained in the EPFL animal facility. To induce pulmonary tumors in KP mice, a lentiviral vector encoding a Cre recombinase was intratracheally delivered to the lungs of 15-week-old mice, as previously described^2,3^. Mice carrying orthotopic KP tumors were generated by intravenously injecting KP cancer cells into 8-week-old wild-type C57BL/6 mice, as previously described^3^. All procedures were conducted according to protocols approved by the Veterinary Authorities of the Canton of Vaud, in obedience to the Swiss laws (protocols 3785a, 3785b, 3785c, 3785d, 3785e, 3785f, 3785g).

### Micro-CT imaging

Lung micro-CT imaging was performed using the Quantum FX micro-CT scanner (PerkinElmer) or SKYSCAN 1276 In Vivo MicroCT (Bruker), with mice anesthetized under continuous isoflurane throughout the procedure. For GEMM KP mice, initial scans confirming tumor establishment were conducted at 11 or 12 weeks after intratracheal administration of the Cre-expressing lentiviral vector. For orthotopic KP tumor-bearing mice, scans were conducted once weekly starting from day 10 or 11 post-cancer cell injection. Imaging was performed weekly until the end of the experiment.

### Manual tumor characterization

To provide tumor annotation data for initial model training and evaluation (see Dataset description), manual tumor annotation was performed using Analyze 12.0 (AnalyzeDirect) for the KP GEMM or OsiriX MD (Pixmeo) for the orthotopic KP model, following previously established standards for identifying murine and human lung tumors in CT scans^3^. To track tumor progression, annotations from different scan time points were aligned to identify corresponding tumors across images. For each matched tumor, both its label and volume were recorded. Fold change (FC) in individual tumor volume or total tumor burden per mouse (sum of its tumor volumes) was calculated using the following formula:

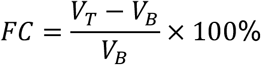

where *V_T_* represents the tumor volume or burden at the evaluated timepoint, and *V_B_* is the baseline value obtained from the initial scan or the last scan prior to treatment.

### Dataset description

To train and evaluate a nnU-Net model for tumor segmentation, we assembled a dataset comprising 1287 scans, originating from both an earlier study^3^ and unpublished experiments. This dataset included 1068 scans from unpublished experiments accumulated for training and 219 scans from the earlier study that were set aside for evaluation. These scans were acquired with a voxel size of 41 μm, and they covered a broad range of tumor sizes and disease stages representative of the diversity of data emanating from preclinical studies in orthotopic and GEMM KP models. To demonstrate the LuTuFlow pipeline, we used CT scan data from two experimental cohorts comprising 8 orthotopic KP mice and 5 GEMM KP mice.

### Scan image processing

Raw data exported from the CT scanner were processed for each mouse to extract 2D TIFF images, which were subsequently stacked into a continuous 3D TIFF volume. The resulting volumetric images could be saved locally or uploaded to an online server for centralized data storage and access, depending on users’ needs. This process is integrated into the Napari-based UI (described below), but it can also be done independently through a Groovy script executed in ImageJ.

### Lung segmentation

The lung region of interest (ROI) was first identified and used to crop the original CT volumes. Working with ROIs reduces data size, accelerates downstream processing, and enables the concatenation of ROIs from multiple consecutive scans to form 4D (3D+time) image series for each mouse. To obtain this ROI, we fine-tuned a pretrained YOLOv8 model (yolov8n-seg) from the Ultralytics Python library^17^ for the task of segmenting lung contours on axial 2D slices. A total of 500 transverse image slices were randomly sampled from the dataset, manually annotated in Napari, and saved in YOLO format. The model was trained on these annotations at the default image size (imgsz) of 640 pixels, for 1,000 epochs, with an 80/20 split between training and validation fractions, and the model checkpoint yielding the best validation performance was retained.

To extract a segmentation mask of the lungs in 3D, the trained model processes a CT scan image slice-by-slice along its transverse axial direction. Each 2D binary segmentation mask predicted for a slice is further refined by morphological hole filling, and only the largest connected component is retained. The 2D masks are subsequently aggregated into a 3D volume, which undergoes two iterations of morphological dilation using a 3D structuring element to compensate for potentially missing slices. The largest connected component in the resulting 3D volume is retained as the final lung segmentation. The 3D bounding box of this segmentation mask is then used to crop the original CT volume.

### Tumor segmentation

Tumor segmentation was performed using the nnU-Net framework^19^ in its 3D full-resolution (3d_fullres) configuration. The model was trained on the extracted lung ROI volumes using a mixture of manually annotated tumors and corrected masks from previous model predictions (see Human-in-the-loop model improvement) as ground truth. 80% of the data was used for training, and 20% was reserved for validation.

For prediction, images are expected to be at a 41 μm resolution, matching the voxel size of the training set images. In preprocessing, voxel intensities (originally in Hounsfield units) are rescaled to the range [0, 1] using min-max normalization. Then, for each 3D input volume, the network generates a binary tumor mask. Individual tumor nodules are delineated by applying connected-component labeling to this binary mask. Tumor nodules smaller than 150 voxels are removed to eliminate spurious detections. Finally, a hole-filling operation is applied to ensure that the segmented regions do not contain any holes. In the resulting cleaned label map, each tumor nodule is assigned a unique identifier, which is subsequently tracked across longitudinal scans to identify individual lesions and their progression.

### Tumor tracking

For the longitudinal tracking of individual tumor nodules, i.e., the process of matching individual tumors over time in order to report their volume change, we start by concatenating all the CT ROIs of a mouse into a single 4D image array, and we do the same with the corresponding tumor masks. Then, we segment the lungs from each scan using a YOLO26^22^ semantic segmentation model (yolo26n-sem), which we fine-tuned specifically for the task of lung segmentation from ROI images. In our pipeline, this process differs from the initial lung detection, which relies on the broader context of the original scans, and is used only to identify the boundaries of the regions of interest. Here, the YOLO model performs a binary semantic segmentation optimized specifically to detect lung boundaries at the scale of the region of interest. This allows us to first align the lung masks, and to use the corresponding spatial transformations to align the tumor masks, which facilitates tumor tracking. More specifically, we reconstruct a surface mesh for each lung mask in the time series using the marching cubes algorithm, then align the vertices of these meshes using the Iterative Closest Point (ICP) method implemented in the Vedo library^23^. We retain the resulting 3D affine transformations and apply them to the tumor masks using nearest-neighbor interpolation. This results in a 4D stack of tumor masks aligned spatially according to the position and deformation of the lungs.

Subsequently, tumor tracking was performed using the “track” function from the LapTrack library^20^. Object merging and splitting were disabled to satisfy the assumption that tumor nodules do not merge or split but remain compact and separate objects throughout the time series. Moreover, a custom cost function was defined to account for both the spatial proximity between tumors in consecutive scans and the relative changes in their volumes. Candidate matches were restricted to objects within a 30-pixel search radius and were accepted only when the relative volume difference did not exceed 100% (a tumor cannot shrink by more than half, or grow to more than double its original size between two consecutive scans). These constraints, combined with the initial spatial alignment, were designed to reduce tracking ambiguities and improve the reliability of automated tumor matching.

The automated tracking module has been implemented in the LuTuFlow pipeline, and is undergoing further optimization and validation. For the longitudinal analyses presented in this study, tumor matching was performed manually by expert investigators (see Manual tumor characterization).

### Software package and user interface

We developed a software package (see **Code availability**) based on Python that includes a Napari^21^ plugin allowing users to load CT images, detect regions of interest, segment tumors, assemble 3D+time series, track tumors, and visualize the results in both 2D and 3D views. Tumor volumes and the associated tracking data can be exported for downstream biomedical analyses. In addition to this graphical user interface, our software package offers a command-line interface (CLI) that allows users to execute the various steps of the workflow directly from the terminal.

### Human-in-the-loop model improvement

To evaluate and improve the accuracy and reliability of the workflow, model-generated tumor annotations were reviewed by biomedical experts. Within the Napari UI, tumor masks were overlaid on the ROI-cropped CT volumes and manually refined to correct imperfect annotations, remove false positives, and recover missed detections. This interactive review process enabled efficient inspection, adjustment, and validation of pipeline-generated results. The curated annotations were subsequently incorporated into the next training iteration of the tumor segmentation model, following the training strategy described above.

## Results

### LuTuFlow streamlines AI-assisted tumor prediction, expert correction, and longitudinal analysis

To streamline scan analysis and reduce subjective bias, we developed an integrated workflow that combines CT image processing, model training, AI-assisted tumor prediction, manual correction, longitudinal tumor tracking, and data export within a single analysis pipeline (**Fig. 1A**). The tumor prediction model was initially trained on manual annotations and subsequently refined using expert-corrected model-generated annotations, establishing a recursive training strategy in which expert feedback improves both output data quality and subsequent model performance. The Napari-based analysis interface provides intuitive access to individual workflow functions, enabling users to interactively inspect, correct, track, and export tumor annotations (**Fig. 1B**).

**Figure 1.**
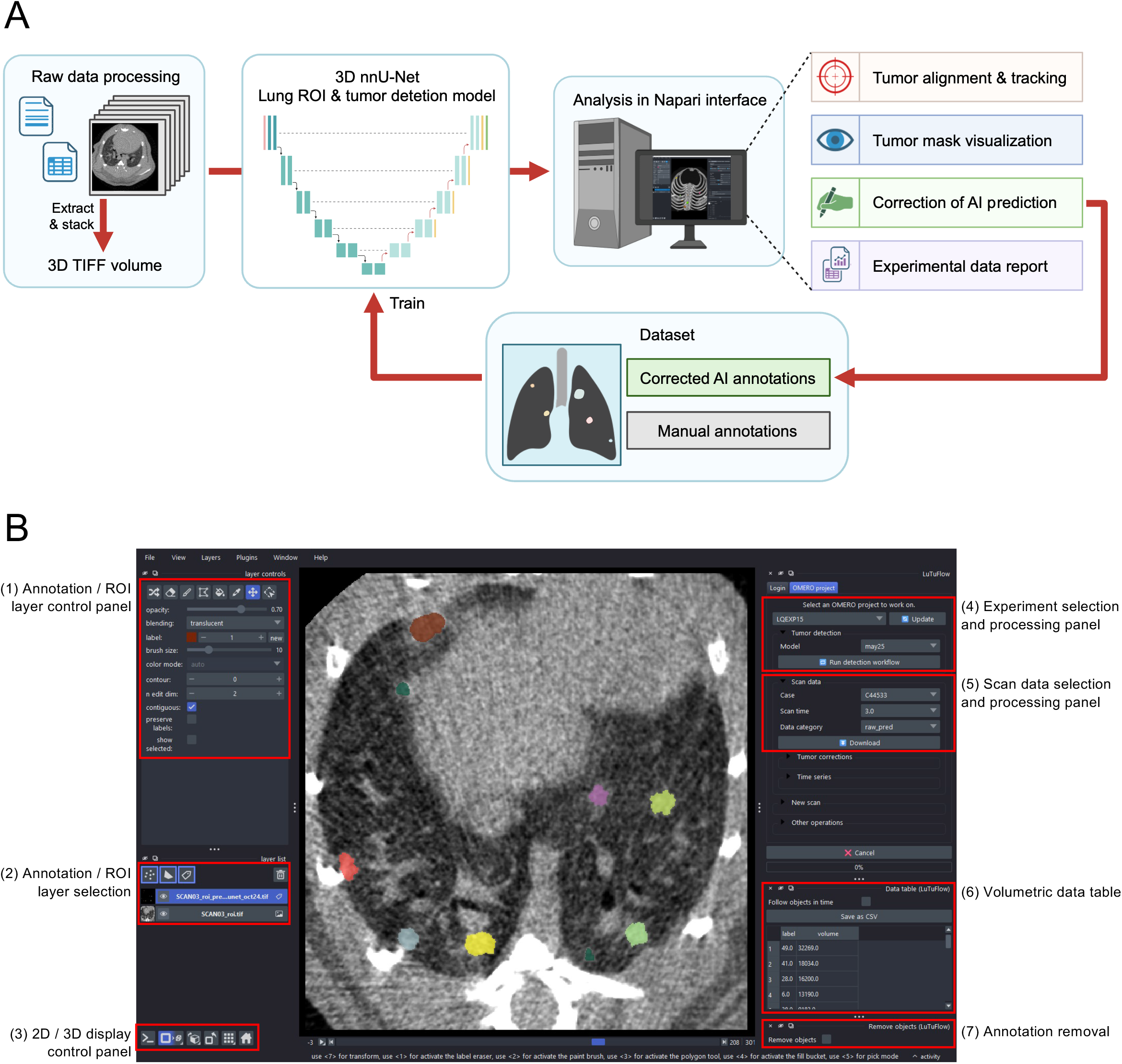
LuTuFlow provides an integrated workflow and user interface for quantitative tumor characterization. **(A)** Overview of the LuTuFlow pipeline. Raw CT scan files are processed by extracting 2D sectional images and reconstructing them into 3D TIFF volumes, which are then used by the lung ROI detection and tumor prediction models. Prediction results are analyzed in the Napari-based interface for tumor visualization, scan alignment, tumor tracking, AI prediction correction, and data export. Corrected annotations can be incorporated into the existing training dataset to recursively improve the model. Red arrows indicate the core procedures in the workflow. **(B)** User interface showing annotated tumor masks overlaid on the scan ROI. (1) Annotation/ROI layer control panel for adjusting image/mask display and editing tumor masks. (2) Annotation/ROI layer selector for switching between scan image and mask layers. (3) 2D/3D visualization control panel for viewing images in different anatomical orientations or as 3D volumes. (4) Experiment-level control panel for uploading and downloading cohort data and initiating automated ROI segmentation and tumor annotation. (5) Scan-level control panel for selecting, uploading, and downloading individual mouse scans and tumor masks. (6) Volumetric data table showing tumor labels and volume measurements in voxels. (7) Object removal tool for deleting individual tumor annotation masks. A detailed user guide is provided in the project GitHub repository.

### LuTuFlow enables longitudinal tumor segmentation and quantification of pulmonary tumor burden

To demonstrate the application of LuTuFlow for longitudinal tumor monitoring, we analyzed CT images from an experiment involving orthotopic KP tumors. In this study, wild-type mice were inoculated with KP cancer cells and subjected to weekly pulmonary CT imaging for three consecutive weeks. LuTuFlow was able to annotate tumors across different stages of tumor growth, including early-stage lesions at week 1 and progressing tumors at week 3, displayed as colored masks overlaid on the 2D CT images or reconstructed 3D CT volumes in the Napari UI (**Fig. 2A**).

**Figure 2.**
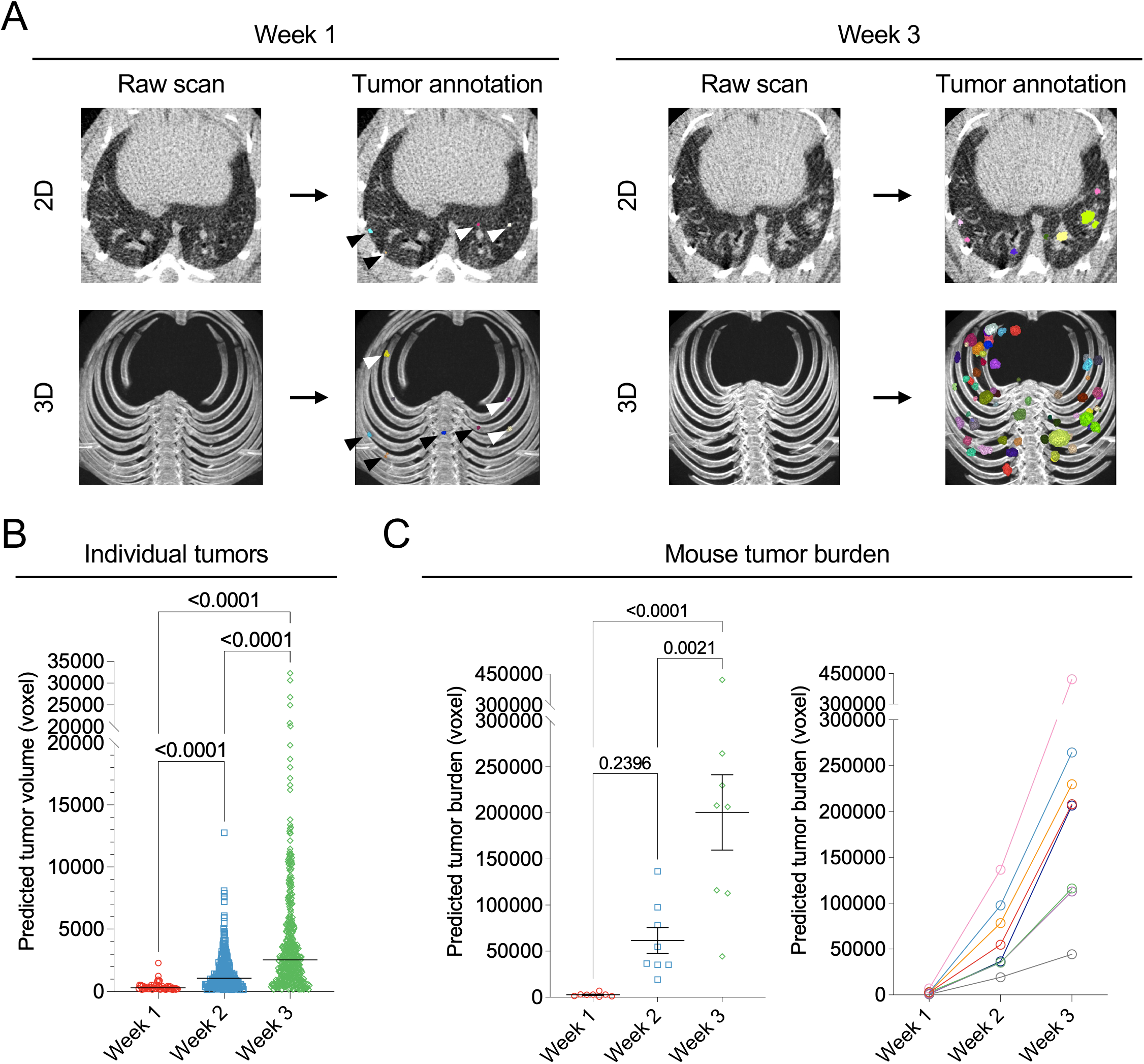
Tumor mask prediction and volume quantification generated by LuTuFlow. **(A)** Representative raw CT images and predicted tumor masks from the same mouse at week 1 and week 3, shown as 2D sections and 3D whole-lung reconstructions. Small nodules in the week 1 annotations are indicated by black and white arrowheads. **(B)** Predicted volumes of individual tumors pooled across all mice at weeks 1, 2, and 3. **(C)** Longitudinal progression of tumor burden in individual mice, calculated as the summed volume of all predicted tumors per mouse. Left panel: cohort summary; right panel: individual-mouse trajectories. Tumor size is presented as voxel counts. Experimental cohort: orthotopic KP tumor model, n = 8 mice, three scans per mouse.

Tumor volumes expressed as voxel counts were then calculated from the 3D volumetric reconstructions of the annotated tumor masks. At the level of individual tumors, pooled analysis of all predicted tumors across mice showed a progressive increase in tumor volume from week 1 to week 3 (**Fig. 2B**). This analysis captures the distribution of tumor sizes within the cohort and highlights the expansion of individual lesions during disease progression.

To gain insight into disease progression at the individual-mouse level, we calculated tumor burden for each mouse as the sum of predicted volumes of all tumors within its lung. This analysis revealed an increase in tumor burden over time in each mouse (**Fig. 2C**); however, when represented longitudinally, progression trajectories differed in individual mice (**Fig. 2C**), reflecting tumor heterogeneity in size and progression rate, as is expected in the KP model^24^. In summary, LuTuFlow can convert serial CT scans into quantitative readouts of tumor burden at both individual tumor and mouse levels.

### Expert validation and correction of AI-generated tumor predictions

To quantitatively evaluate model performance and provide curated annotations for subsequent training iterations, AI-generated tumor annotations from week 1 and week 3 scans were examined and corrected by expert investigators. The manually corrected annotations were used as the reference ground truth, against which raw model predictions were compared. Correction events were categorized as true positives (TP), false positives (FP), or false negatives (FN). TP annotations correspond to predicted masks located at the correct tumor position, including those with accurate tumor coverage as well as those with moderate discrepancies in shape or volume. The latter were manually refined to better match the tumor boundary (**Fig. 3A**). FP annotations represented model detections that did not correspond to true tumors, including structures outside the lung parenchyma or cavities within bronchi and alveoli. FPs were manually removed during correction (**Fig. 3B**). FN events corresponded to tumors missed by the model and were manually added by expert investigators (**Fig. 3C**). Across all evaluated objects, including predicted tumors and missed lesions, TP annotations accounted for 75.26% of the total events, representing the majority of correction outcomes (**Fig. 3D**).

**Figure 3.**
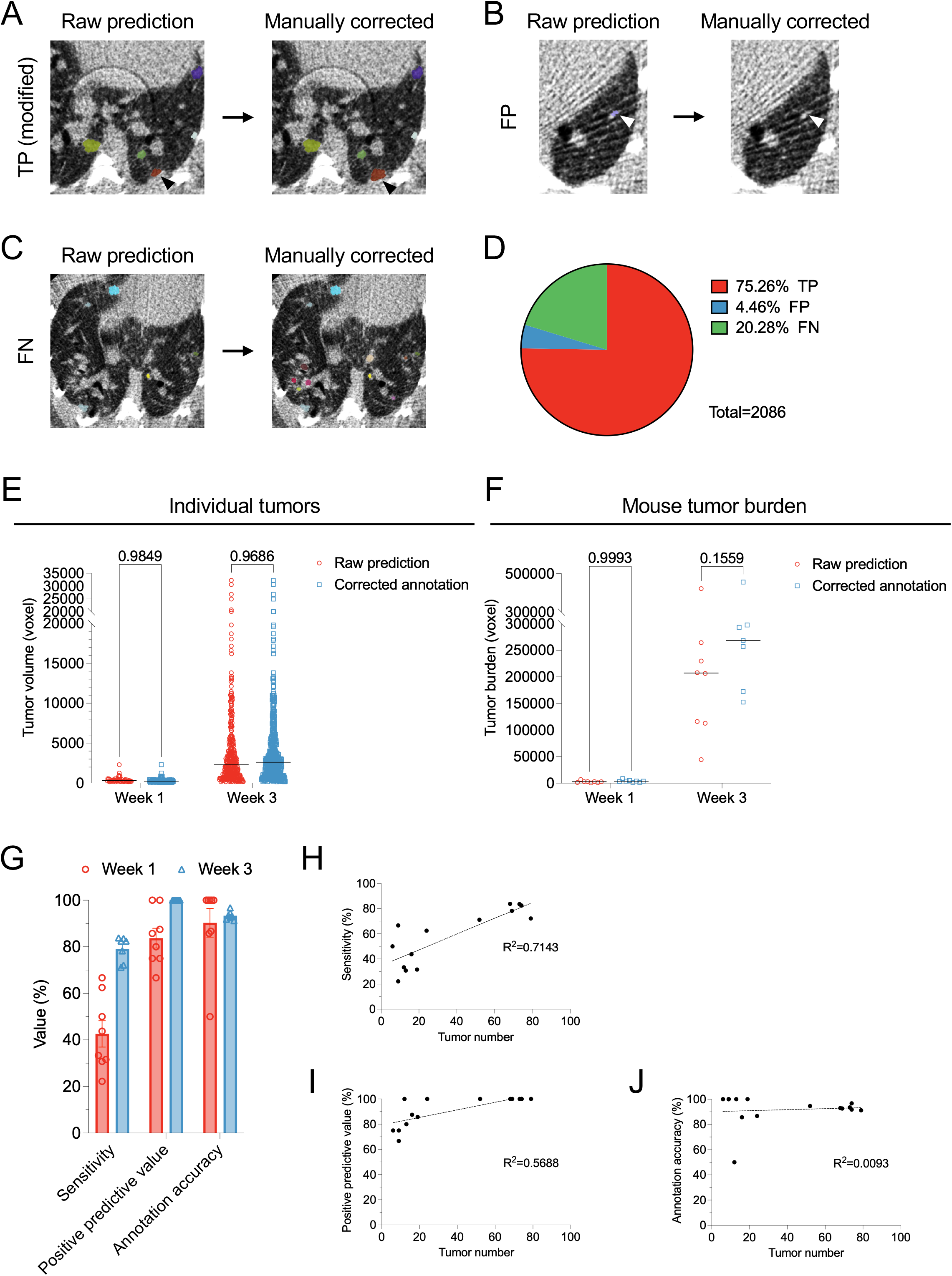
Correction of model-generated tumor annotations and evaluation of model performance. **(A)** Example of a true positive (TP) tumor annotation modified during manual correction, indicated by an orange lesion and a black arrowhead. **(B)** Example of a false positive (FP) annotation removed during manual correction, indicated by a white arrow. **(C)** Example of false negative (FN) tumors added during manual inspection. **(D)** Pie chart showing the distribution of TP, FP, and FN events across all evaluated scans. **(E)** Comparison of individual tumor volumes exported from raw predictions and manually corrected annotations at week 1 and week 3. **(F)** Comparison of mouse-level tumor burden calculated from raw predictions and manually corrected annotations at week 1 and week 3. **(G)** Model performance metrics, including sensitivity, positive predictive value, and annotation accuracy. Each point represents the value calculated from tumors within each mouse. (**H**-**J**) Correlations between model performance metrics and tumor number, with week 1 and week 3 tumors pooled for analysis. Tumor size is presented as voxel counts. The experimental cohort is the same as in Figure 2.

We then assessed whether expert correction substantially altered tumor quantification. Tumor progression was compared between raw model predictions and manually corrected annotations at both individual tumor and mouse levels. When individual tumor volumes were pooled across mice, raw predictions and corrected annotations showed no statistically significant differences at either week 1 or week 3 (**Fig. 3E**). A similar pattern was observed for mouse-level tumor burden (**Fig. 3F**). Corrected annotations at week 3 showed a tendency toward higher tumor burden compared with raw predictions, but the difference remained non-significant. These results indicate that raw and corrected annotations yielded similar cohort-level estimates, supporting reliable quantitative agreement between model predictions and expert-corrected annotations.

To further characterize model performance, we calculated three complementary metrics referring to established standards for diagnostic testing^25^: sensitivity, positive predictive value, and annotation accuracy (**Fig. 3G**). Sensitivity was defined as the number of TP tumors divided by the total number of real tumors after correction, namely TP/(TP+FN), reflecting the ability of the model to identify tumors. Positive predictive value was calculated as the number of TP predictions divided by all model-predicted tumors, i.e., TP/(TP+FP), indicating the fraction of predicted lesions that corresponded to true tumors. Annotation accuracy was defined as the fraction of TP tumors that required no manual modification, reflecting the ability of the model to correctly annotate detected tumors. Among these metrics, sensitivity was lowest at week 1, indicating that the model was less effective in detecting early-stage disease when there were fewer and smaller tumors. Consistent with this observation, sensitivity showed a positive association with tumor number, suggesting reduced detection sensitivity in scans with lower tumor numbers (**Fig. 3H**). In contrast, positive predictive value and annotation accuracy were generally higher and showed no clear association with tumor number (**Fig. 3I,J**). Together, these analyses suggest that LuTuFlow produced accurate predictions once tumors were detected, whereas the main limitation of the model was reduced sensitivity for very small or sparse lesions at disease onset.

### LuTuFlow facilitates monitoring of the tumor progression

We next used the Napari-based UI of LuTuFlow to precisely track tumor progression at the level of individual lesions. For this analysis, we used CT scans from an independent cohort of 5 GEMM KP mice, which were followed for four weeks with weekly pulmonary CT imaging (weeks 1-4). In this model, fewer lung tumors develop in each mouse than in the orthotopic KP model^3^. Predicted tumor annotations were manually corrected as described above. Within the Napari interface, serial scans from the same mouse were displayed side by side to identify tumors that could be matched across timepoints (**Fig. 4A**). For each tracked tumor, the corresponding label and volume were recorded at each scan.

**Figure 4.**
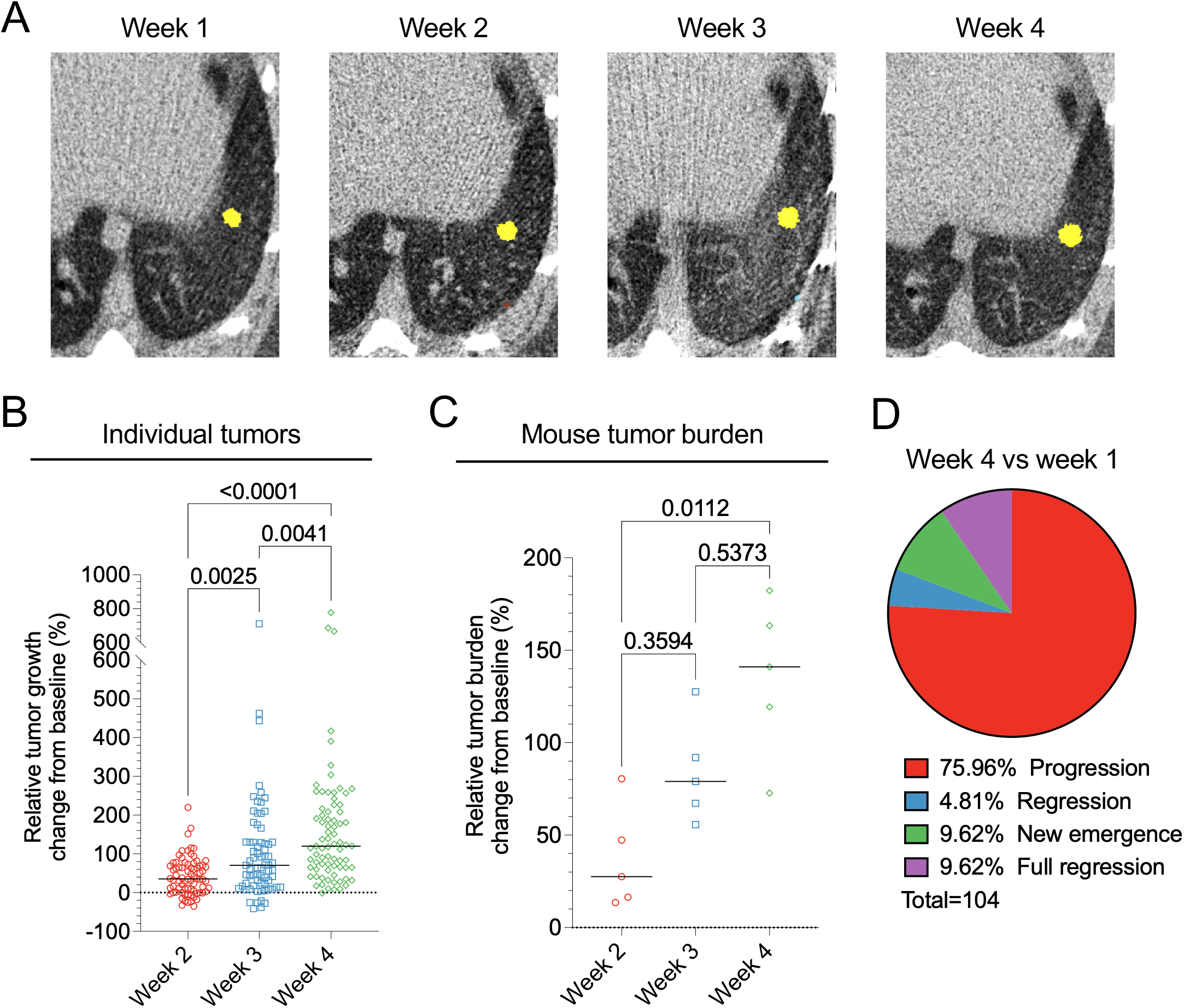
Demonstration of longitudinal tumor tracking and tumor progression classification. **(A)** Representative example of a matched tumor (pseudocolored in yellow) from the same mouse across serial scan time points from week 1 to week 4. **(B)** Growth of individual tracked tumors at weeks 2, 3, and 4, relative to the week 1 baseline. **(C)** Mouse-level tumor burden progression at weeks 2, 3, and 4, relative to the week 1 baseline. **(D)** Pie chart showing the distribution of tumor progression events at week 4 compared with week 1, including progression, regression, new emergence, and full regression. Experimental cohort: GEMM KP, n = 5 mice, four scans per mouse.

Using week 1 as the baseline, we calculated the relative growth of each individual tumor by dividing the volume change at later time points during the experiment by its baseline volume (see **Methods**). This enabled lesion-level assessment of tumor progression across longitudinal scans (**Fig. 4B**). In parallel, mouse-level tumor progression was quantified by calculating the relative change in total tumor burden for each mouse compared with the week 1 baseline (**Fig. 4C**). Both lesion-level and mouse-level analyses revealed an overall increase in tumor growth over time, with variable progression patterns across tumors and mice, as expected in this GEMM^3^.

Tracking tumors at the individual-lesion level provides information beyond total tumor size or whole-mouse tumor burden. During disease progression, some small lesions detected at earlier time points were no longer detectable at later scans, whereas new tumors can emerge over time. As a result, not all lesions can be definitively matched across the full longitudinal series. We therefore classified tumor dynamics into progression, regression, new emergence, and full regression (**Fig. 4D**). This lesion-level analysis may be useful for evaluating tumor progression during therapeutic intervention, where regression of individual lesions occurs more frequently than in untreated mice^3^.

## Discussion

Quantitative evaluation of tumor growth is critical for preclinical studies of pulmonary cancer, whereas conventional manual tumor annotation is labor-intensive and prone to inconsistency^6,7^. Here, we developed LuTuFlow, an integrated pipeline for AI-assisted analysis of longitudinal pulmonary CT scans in mouse tumor models. By combining automated segmentation, tumor burden quantification, expert-guided correction, iterative model training, and longitudinal tracking, LuTuFlow provides a practical tool for analyzing CT imaging data for quantitative readouts of lung tumor progression.

LuTuFlow facilitates the processing of scan images into tumor volume measurements at both individual-lesion and whole-mouse levels. In our demonstration, both pooled lesion-level analysis and mouse-level tumor burden reliably captured progressive tumor growth over time. These complementary readouts are useful because different preclinical settings may require different analytical levels. Mouse-level tumor burden provides an integrated measure of disease progression, whereas lesion-level analysis reveals tumor heterogeneity and individual growth behavior.

Manual correction did not substantially alter cohort-level tumor quantification, suggesting that raw predictions were sufficient to reflect overall longitudinal tumor progression in the demonstrated cohort. Nevertheless, expert correction remains important for quality control, performance assessment, and recursive model refinement. Corrected annotations provide curated training data for subsequent training cycles, allowing the model to improve as additional datasets become available. In our case, retraining improved model performance, although further gains may reach a plateau as the dataset grows and the model approaches the performance required for preclinical applications. The iterative training strategy also allows incorporation of data from different tumor models, supporting adaptation to diverse preclinical applications.

Model performance was strongly influenced by raw scan quality, which affected the detection of small lesions. High-contrast anatomical structures, including bronchi and bones, or high background noise can obscure tumor boundaries and reduce contrast between small tumors and surrounding lung tissue. These limitations are especially relevant for early-stage lesions, which are often small, sparse, and susceptible to partial-volume effects, low contrast, and confusion with adjacent anatomical structures. Accordingly, the main limitation of the current model was reduced sensitivity for early-stage lesions, consistent with previous reports showing that image-based tumor delineation is influenced by lesion size, imaging heterogeneity, and segmentation method^26^. However, this limitation had little impact on volumetric analysis, as missed early lesions contributed minimally to total tumor burden. This is consistent with the lack of a significant difference between raw predictions and manually corrected annotations when analyzed as pooled individual tumor volumes or mouse-level tumor burden.

At later disease stages, manually corrected annotations tended to yield higher tumor volumes, particularly at the mouse-burden level. This likely reflects conservative predicted masks that required manual refinement to fully cover larger or merged tumor regions, consistent with annotation accuracy not reaching 100% in individual mice. These results highlight a potential trade-off between detection sensitivity and the reliability of positive predictions, as reflected by the positive predictive value. Increasing sensitivity for very small lesions may reduce missed tumors but also increase false positives, particularly where small nodules resemble anatomical structures or imaging artifacts. For longitudinal preclinical studies, the most relevant readouts are often total tumor burden and growth trajectories rather than exhaustive counting of very small lesions. Therefore, while improving early-lesion sensitivity remains important, the current model already provides reliable quantitative readouts for monitoring overall tumor progression. Moreover, although longitudinal tracking extends LuTuFlow beyond tumor burden measurement by evaluating progression at the level of individual nodules, fold change cannot be calculated for newly emerged or fully regressed tumors, limiting purely quantitative representation of these events.

A strength of LuTuFlow is its integration into an interactive Napari-based UI. Rather than relying on separate manual viewers or standalone scripts, the workflow allows users to visualize CT volumes, inspect segmentation masks, correct annotations, export quantitative data, and perform longitudinal analysis in a unified environment. This reduces the technical barrier for biomedical researchers without extensive programming experience while preserving flexibility for expert review and model refinement. Compared with purely manual ROI-based viewers, AI-assisted prediction with interactive correction provides a more scalable approach for preclinical lung tumor analysis.

Although the Napari UI enables side-by-side inspection of scans from different time points and facilitates identification of corresponding tumors, manual inspection and recording of tumor matches are still required. Further development will focus on improving and validating automated tracking, especially for newly emerging lesions, disappearing tumors, or spatially adjacent nodules. Another future direction is adapting LuTuFlow to additional preclinical tumor models and, potentially, clinical settings using repeated volumetric scans to monitor disease progression or therapeutic response. Extension to other disease models or species will require model retraining and input from domain experts.

In summary, LuTuFlow provides an integrated and automated CT analysis pipeline for longitudinal tumor monitoring in mouse models. The model generates reliable tumor predictions for cohort-level tumor progression analysis, while expert-guided correction improves annotation quality and supports recursive model refinement. Although sensitivity for small early lesions and automated tracking accuracy remain areas for improvement, LuTuFlow establishes a scalable platform for quantitative, interactive, and longitudinal analysis of preclinical pulmonary tumor models.

## Author contributions

Y.L., L.L. and A.M.-U. conceptualized the LuTuFlow workflow, designed the study, performed experiments, generated corrected annotation data, analyzed and interpreted the results, and wrote the manuscript. M.W. developed the computational methodology, trained the model, implemented and optimized the software, curated the data, analyzed the results, and contributed to writing of the manuscript. N.M. critically reviewed CT scans and tumor annotation data. B.D. contributed to software development. E.A. and M.D.P. coordinated and supervised the study, interpreted the results, and contributed to writing of the manuscript.

## Acknowledgements

The authors thank Alan Guichard, Bruno Torchia and Celine Wyser-Rmili for managing mouse colonies and acknowledge the excellent technical support of the scientific platforms at EPFL and the Agora Cancer Research Center. This study was supported by grants from a generous donor advised by Carigest SA (to M.D.P.), the ISREC Foundation (Tandem to M.D.P. and N.M.), the Fondation Chercher et Trouver via Swiss Cancer Research (KFS-6045-02-2024 to M.D.P.), and the European Research Council (ERC-CoG EVOLVE-725051 to M.D.P.).

## Code availability

All the source code and distributions of our pipeline can be found in our GitHub repository. The user interface to access all the mentioned functions in our pipeline can be found at: https://github.com/EPFL-Center-for-Imaging/lutuflow.

Model weights from different iterations of the tumor prediction model have been deposited at: https://sandbox.zenodo.org/records/588733.

## Declaration of generative AI in the writing process

During the writing of this manuscript, some of the authors used ChatGPT to improve the language and readability of the text. The authors reviewed and edited the content as necessary after using this tool and take full responsibility for the content of the published manuscript.

